# Substrate-specific regulation of the mTORC1 pathway by G protein-coupled receptors

**DOI:** 10.1101/2024.09.18.613687

**Authors:** Samuel J. Atkinson, Florentina Negoita, William Vincent Ritchie, Kyle Thompson, Kei Sakamoto, Dawn Thompson, James N. Hislop, Riko Hatakeyama

## Abstract

The mammalian/mechanistic Target of Rapamycin Complex 1 (mTORC1) orchestrates cell growth and metabolism in response to diverse external cues. mTORC1 has a myriad of phosphorylation substrates, each playing important physiological roles. Emerging evidence suggests that mTORC1 can respond to upstream signals in a nuanced manner, enabling differential regulation of individual substrates and downstream biological processes. However, the nature of signals that determine the signaling selectivity of mTORC1 remains incompletely understood. Here, we studied mTORC1 regulation by G protein-coupled receptors (GPCRs). We found that phosphorylation of the Transcription Factor EB (TFEB), a non-canonical mTORC1 substrate that controls lysosome biogenesis, responds to GPCRs differently, compared to canonical mTORC1 substrates controlling protein synthesis such as S6K1 and 4EBP1. In particular, the muscarinic acetylcholine receptor M5 (M5R) promoted phosphorylation of S6K1 and 4EBP1, while triggering TFEB dephosphorylation. Consequently, M5R stimulated protein synthesis without inhibiting lysosome biogenesis. The regulations of an anabolic process and a catabolic process, albeit both governed by mTORC1, are thus decoupled downstream of M5R. This study highlights the importance of reassessing the effects of GPCRs on mTORC1 by concurrently monitoring individual substrates, a critical consideration to be made when evaluating GPCR ligands as therapeutic agents targeting the mTORC1 pathway.

## Introduction

Cells adapt to the changing environment by fine-tuning various biological processes. At the core of the cellular adaptation mechanism is the mammalian/mechanistic Target of Rapamycin Complex 1 (mTORC1), a master regulator of cell growth and metabolism (1)(2). mTORC1 is a serine/threonine kinase that gets activated by pro-growth signals such as nutrients (e.g., amino acids) and growth factors (e.g., insulin). mTORC1, in turn, activates pro-growth cellular processes such as anabolic reactions (e.g. protein synthesis) while repressing catabolic reactions (e.g. protein degradation via autophagy), through phosphorylating its substrate proteins. mTORC1 activation by amino acids and by insulin/growth factors is mediated by Rag and Rheb small GTPases, respectively. mTORC1 plays a pivotal role in cellular/organismal homeostasis in humans and is implicated in a range of conditions, including cancer, diabetes, neurodegeneration, and aging. Developing effective therapeutic strategies for these diseases requires a deeper mechanistic understanding of mTORC1 regulation.

An emerging theme in the mTORC1 research field is mTORC1’s differential signaling to individual substrates. Depending on upstream stimuli, the phosphorylation patterns of several “non-canonical” mTORC1 substrates including the transcription factor EB (TFEB), a regulator of lysosome biogenesis, responds differently to that of canonical mTORC1 substrates such as the ribosomal protein S6 kinase 1 (S6K1) and the eukaryotic translation initiation factor 4E-binding protein 1 (4EBP1), regulators of global protein synthesis (3)(4)(5). Unlike phosphorylation of canonical substrates that requires activation of both Rag and Rheb small GTPases, TFEB phosphorylation is thought to only respond to Rag GTPases.

Two models have been proposed to explain the distinct behavior of TFEB, which are not mutually exclusive. One model attributes the difference to the unique substrate recruitment mechanism, in which Rag GTPases physically interact with TFEB (but not S6K1 or 4EBP1) and present it to mTORC1 (3)(6). Another model assigns these substrates to spatially distinct mTORC1 pools; the cytosolic mTORC1 pool phosphorylates S6K1 and 4EBP1, while the lysosomal mTORC1 pool phosphorylates TFEB (5). In either case, a comprehensive view is missing on how various external cues, that are typically sensed by cell-surface receptors, differentially feed into individual mTORC1 substrates.

G protein-coupled receptors (GPCRs) constitute the largest cell-surface receptor family, being encoded by more than 800 genes in the human genome (7)(8). GPCRs sense diverse signals ranging from chemicals such as growth factors and odorants to physical parameters such as light.

Characterizing any GPCR-mTORC1 signaling therefore has wide-reaching implications for cellular adaptation mechanisms operating in diverse contexts across the human body. Moreover, GPCRs are highly druggable proteins, as more than 30% of FDA-approved drugs target GPCRs (9). To harness the therapeutic potential of these drugs and enable their safe and effective repurposing across diverse diseases, it is essential to elucidate how G protein–coupled receptors (GPCRs) transmit signal to downstream pathways such as the mTORC1 pathway.

For many years GPCR signaling was thought to occur exclusively via activation of heterotrimeric G-proteins, with distinct Gα subtypes triggering distinct downstream events. G_q_ stimulates phospholipase C; G_s_ and G_i_ stimulates and inhibits, respectively, cyclic AMP production; G_12/13_ activates Rho small GTPases (10)(11). This original paradigm has since been challenged, with the identification of G protein-independent signaling cascades, often proposed to be mediated by β-arrestin proteins (12). More recently, the GPCR field has undergone a further paradigm shift. Both G-protein dependent and -independent signaling can occur from intracellular compartments, particularly endosomes, previously thought to be refractory for signaling (13). However, the role of intracellular signaling remains poorly understood, particularly in the context of mTORC1 regulation.

Substantial efforts have been made to elucidate how GPCRs modulate mTORC1 signaling (reviewed in (14)), with some GPCRs shown to activate, while others inhibit, mTORC1. A largely overlooked aspect, however, is the substrate specificity of GPCR-mTORC1 signaling, as most prior studies monitored phosphorylation of only canonical mTORC1 substrates such as S6K1 and/or 4EBP1. In this study, we examined in parallel the response of S6K1, 4EBP1, and the non-canonical mTORC1 substrate TFEB to GPCR stimulation. We found that TFEB responds to GPCR stimulation differently to S6K1 and 4EBP1, allowing simultaneous operation of protein synthesis and lysosome biogenesis. Our findings reveal substrate/downstream specificity as a novel layer of the GPCR-mTORC1 signaling.

## Results

### Development of M5R-DREADD cell line

We examined how phosphorylation of various mTORC1 substrates —S6K1, 4EBP1, and TFEB— responds to GPCR stimulation. As G_q_ has been previously implicated in mTORC1 signaling (14), we investigated a G_q_-coupled GPCR with the muscarinic acetylcholine receptor M5 (M5R) as a model.

Acetylcholine, the endogenous ligand of M5R, stimulates nicotinic and muscarinic receptor subtypes, a number of which are endogenously expressed in HEK293 cells (15) (Figure 1A left). To specifically observe the effects of M5R activation, we developed a system to selectively stimulate M5R. We took advantage of the previously described mutations that make muscarinic receptors into ‘designer receptor exclusively activated by designer drugs (DREADD)’ receptors, which are uniquely activated by the synthetic ligand clozapine N-oxide (CNO) (Figure 1A right) (16).

**Figure 1.**
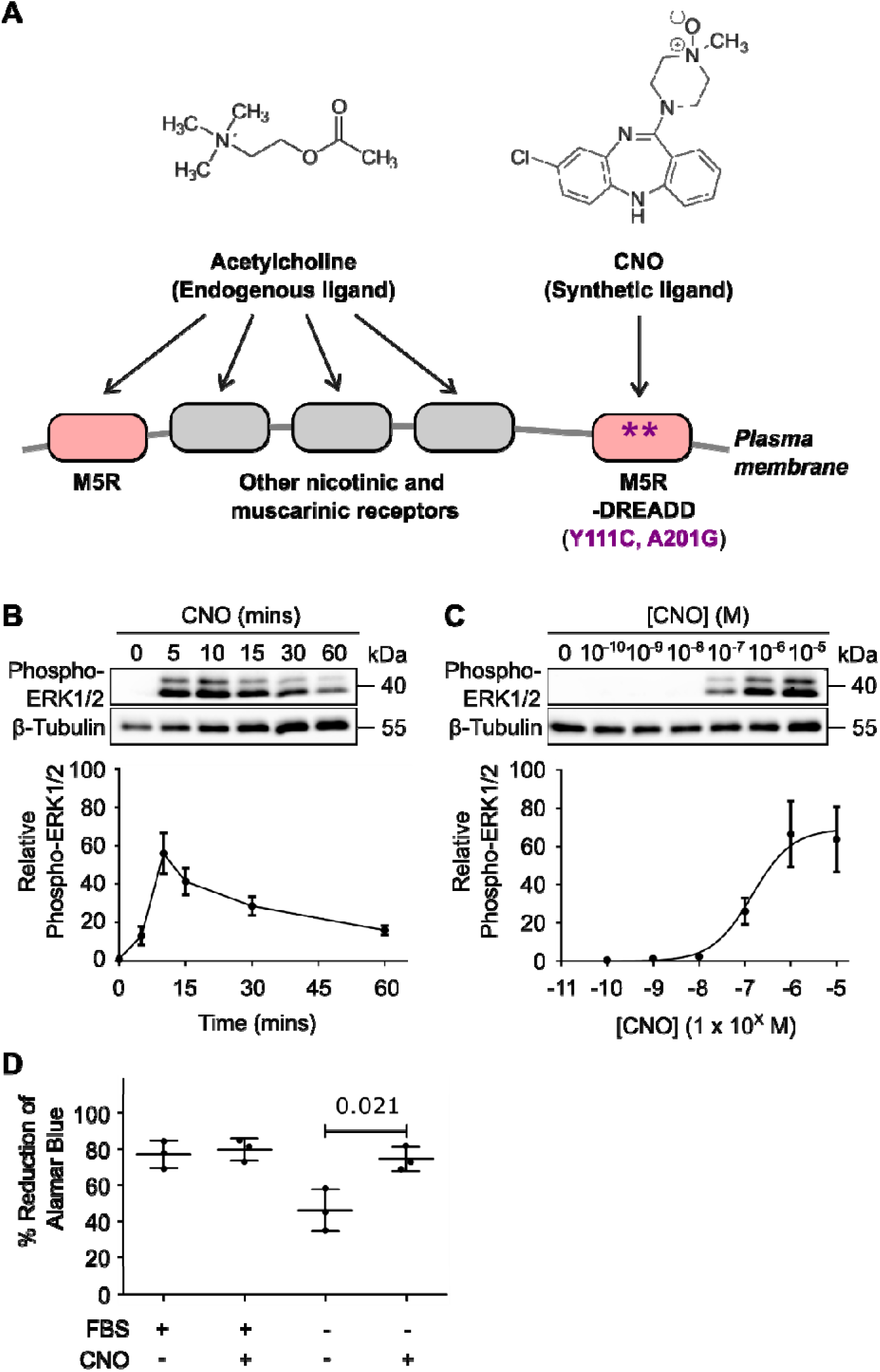
Construction and validation of the M5R-DREADD cell line. (A) Experimental setting for M5R-specific stimulation. M5R was converted into M5R-DREADD by introducing the indicated point mutations. In cells stably expressing M5R-DREADD, the synthetic ligand CNO selectively stimulates M5R-DREADD, unlike the endogenous M5R ligand acetylcholine which binds to nicotinic and muscarinic receptor subtypes. (B) HEK293 cells expressing M5R-DREADD were serum starved for 2 hours prior to stimulation with 1 μM CNO for the indicated time. Cells were lysed and analysed by SDS-PAGE and western blotting. Quantitative analysis of N = 3 experiments is shown at the bottom. (C) HEK293 cells expressing M5R-DREADD were serum starved for 2 hours prior to stimulation with the indicated concentration of CNO for 7 minutes. Cells were lysed and analysed by SDS-PAGE and western blotting. Quantitative analysis of N = 3 experiments is shown at the bottom. (D) Metabolic activity of HEK293 cells expressing M5R-DREADD was measured by the reduction of Alamar Blue after overnight stimulation with 1 µM of CNO with or without serum starvation.

We generated a HEK293 cell line stably expressing M5R-DREADD. The M5R-DREADD cells responded to CNO in a dose-dependent manner as expected, triggering phosphorylation of its downstream MAP kinase ERK1/2 (Figure 1B and 1C, showing the activation kinetics and concentration-response curve, respectively). CNO robustly enhanced the metabolic activity (reducing power) of serum-starved M5R-DREADD cells (Figure 1D), suggesting the positive role of M5R in cell growth and/or proliferation. Our data established the M5R-DREADD cell line as a useful model to investigate growth/proliferation signaling, including the mTORC1 pathway.

### M5R regulates mTORC1 in a substrate-specific manner

We examined phosphorylation of canonical mTORC1 substrates, S6K1 and 4EBP1, that promote global protein synthesis. We serum-starved M5R-DREADD cells to bring the mTORC1 activity down to a basal level before stimulation with CNO. CNO treatment triggered phosphorylation of S6K1, as probed by the phospho-specific antibody (Figure 2A and 2B, panel 1, for a short and long time-course, respectively). This effect of CNO was specifically through M5R stimulation, because it was not seen in the parental wild-type HEK293 line not expressing M5R-DREADD (Figure 2C). Phosphorylation of S6K1 coincided with that of 4EBP1, which was detected as a stereotypical shift in banding pattern to that of higher molecular weight species (Figure 2A and 2B, panel 3).

**Figure 2.**
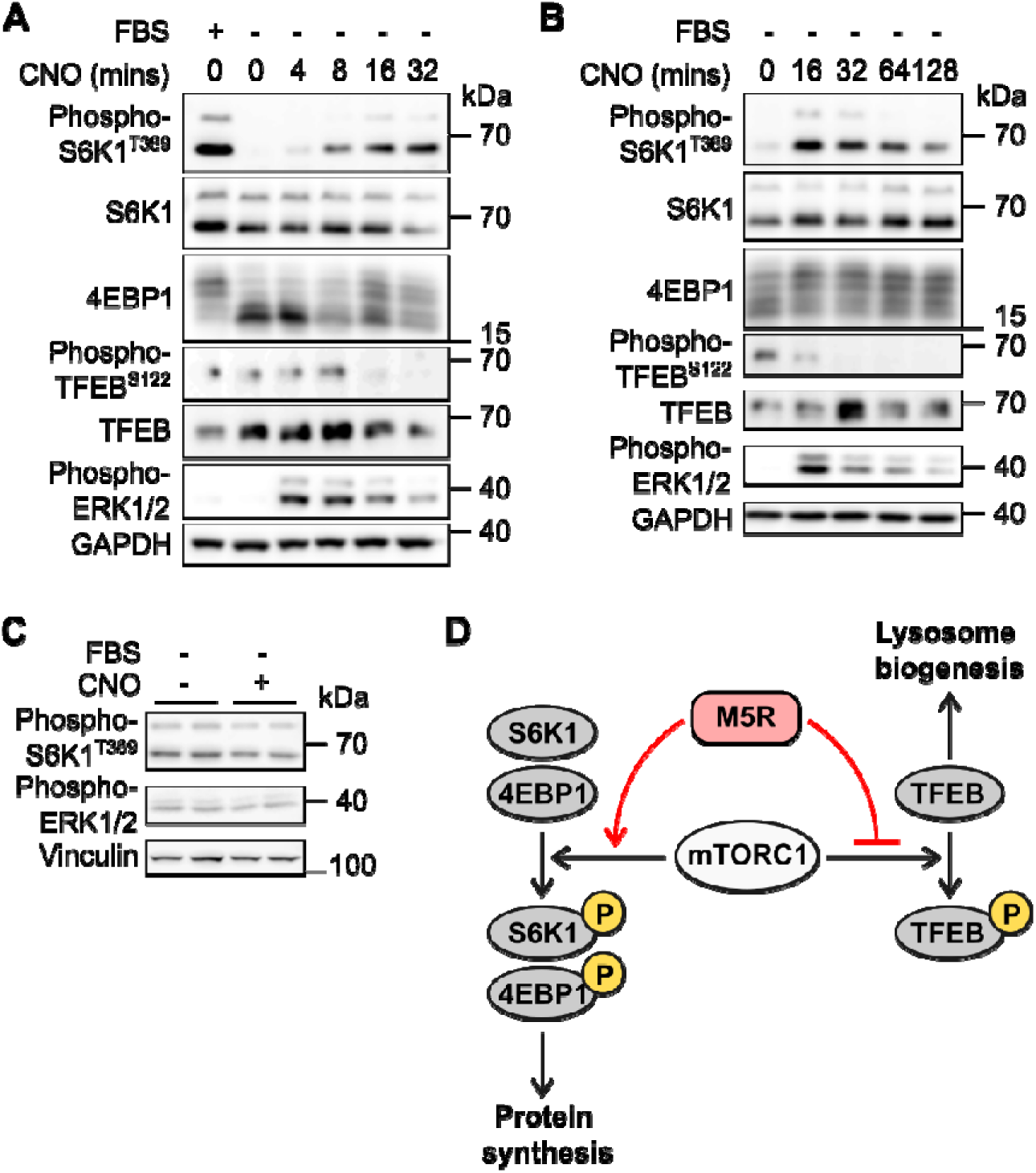
Substrate-specific regulation of mTORC1 by M5R. (A) HEK293 cells expressing M5R-DREADD were starved of FBS or given fresh FBS-containing media for 2 hours before being stimulated with 1 µM CNO for up to 32 minutes. Cells were lysed and analysed by SDS-PAGE and western blotting. (B) HEK293 cells expressing M5R-DREADD were starved of FBS or given fresh FBS-containing media for 2 hours before being stimulated with 1 µM CNO for up to 128 minutes. Cells were lysed and analysed by SDS-PAGE and western blotting. (C) Wild-type HEK293 cells were starved of FBS or given fresh FBS-containing media for 2 hours before being stimulated with 1 µM CNO for 32 minutes. Cells were lysed and analysed by SDS-PAGE and western blotting. (D) Schematic showing the proposed dualistic regulation of mTORC1 substrates by M5R. S6K1/4EBP1 phosphorylation is promoted and TFEB phosphorylation is suppressed by M5R, resulting in simultaneous activation of all three substrates.

We then examined the lysosome biogenesis regulator TFEB, as its phosphorylation by mTORC1 is regulated in a non-canonical, Rheb-independent manner. Strikingly, phosphorylation of TFEB, as probed by the phospho-specific antibody, was rather downregulated by CNO treatment after 16 minutes (Figure 2A and 2B, panel 4). These results suggest that M5R activates mTORC1 towards S6K1 and 4EBP1, while inhibiting it towards TFEB (Figure 2D).

We noticed the slow kinetics of mTORC1 substrate phosphorylation by M5R, which did not peak until 32 minutes in the case of S6K1 (Figure 2A and 2B, panel 1). This was in stark contrast to the kinetics of ERK1/2 phosphorylation that peaked at, or potentially even before, 4 minutes of CNO treatment (Figure 2A, panel 6). This observation raises the intriguing possibility that mTORC1 is activated only after M5R is internalized (see later). M5R appears to transmit qualitatively different signals over time, an early signal via ERK1/2 and a late signal via mTORC1.

### M5R stimulates protein synthesis

Having observed the opposite effects of M5R activation on phosphorylation of canonical substrates (S6K1 and 4EBP1) and that of TFEB, we then examined further downstream cellular processes. Phosphorylation of S6K1 and 4EBP1 by mTORC1 promotes global protein synthesis (1)(2). S6K1 exerts this effect partly through phosphorylation of the ribosomal protein S6. As expected, CNO robustly triggered S6 phosphorylation in M5R-DREADD cells (Figure 3A).

**Figure 3.**
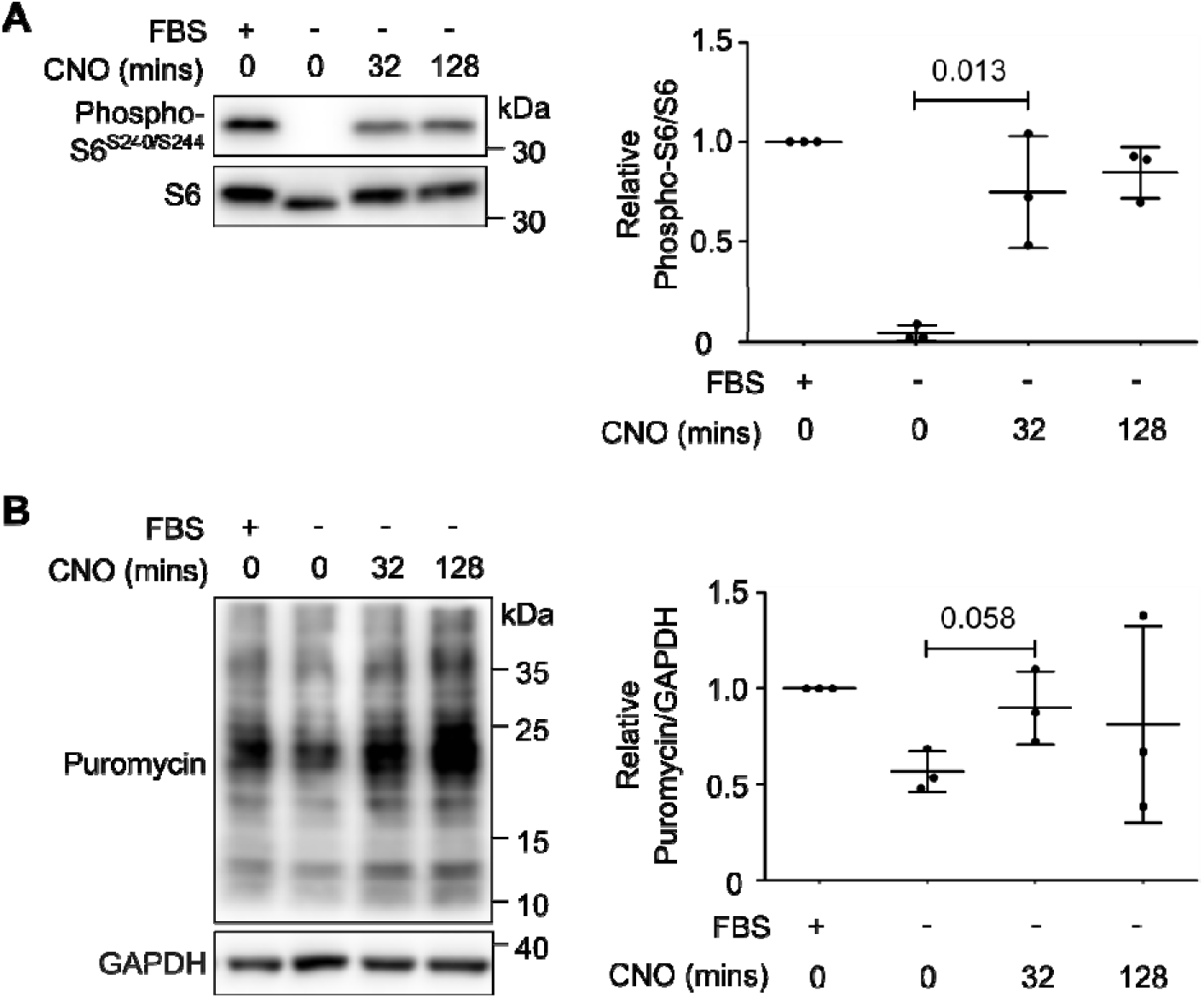
M5R stimulates protein synthesis. (A) HEK293 cells expressing M5R-DREADD were starved of FBS or given fresh FBS-containing media for 2 hours before being stimulated with 1 µM CNO for the indicated times. Cells were lysed and analysed by SDS-PAGE and western blotting. Quantitative analysis of N = 3 experiments is shown on the right. (B) HEK293 cells expressing M5R-DREADD were starved of FBS or given fresh FBS-containing media for 2 hours before being stimulated with 1 µM CNO for the indicated times. Puromycin was added at 5 µg/ml to each condition for 15 minutes before lysis and analysis by SDS-PAGE and western blotting. Quantitative analysis of N = 3 experiments is shown on the right.

*De novo* protein synthesis can be monitored by the established puromycin incorporation assay (17). As shown in Figure 3B, serum starvation caused decreased incorporation of puromycin, suggestive of slowed protein synthesis. Subsequent M5R stimulation by CNO triggered increased puromycin incorporation. These results revealed the positive effect of M5R signaling on cellular protein synthesis activity, consistent with increased phosphorylation of S6K1 and 4EBP1.

### M5R does not inhibit lysosome biogenesis

M5R stimulation caused dephosphorylation of TFEB (Figure 2A and 2B). TFEB dephosphorylation upon amino acid starvation triggers its nuclear translocation, and through activating a transcriptional program, promotes lysosome biogenesis (18). We examined these downstream events upon M5R stimulation.

As expected, CNO treatment in serum-starved M5R-DREADD cells triggered nuclear translocation of TFEB-GFP, similar to amino acid-starved cells (Figure 4A). We confirmed this result by cellular fractionation of cytosol and nuclei. In both serum-fed and -starved cells, more TFEB was detected in the nuclear fraction in CNO-treated samples (Figure 4B, panel 1). The mTOR inhibitor Torin1 similarly caused nuclear translocation of TFEB, as previously reported (18).

**Figure 4.**
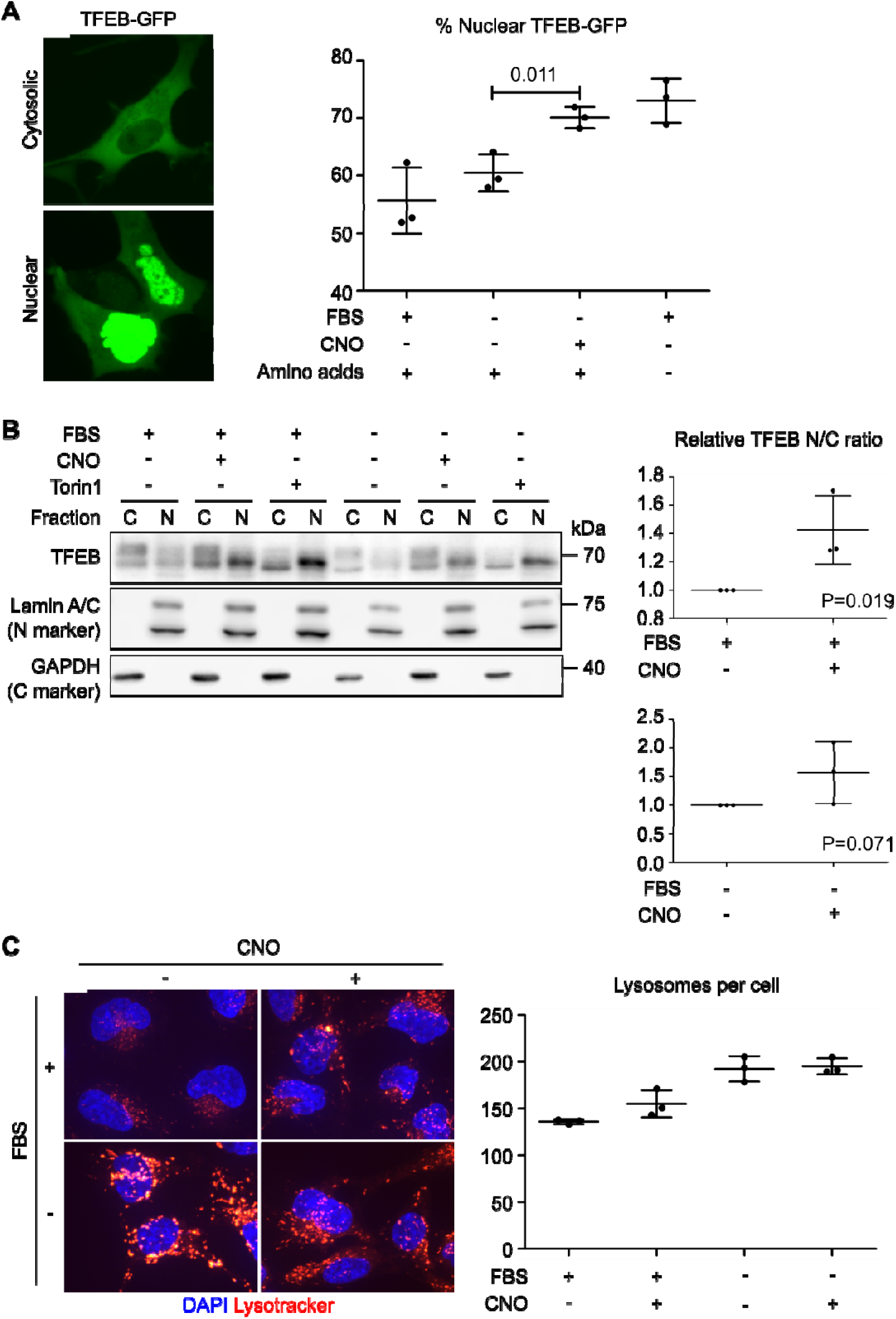
M5R does not inhibit lysosome biogenesis. (A) HEK293 cells expressing M5R-DREADD were transfected with a TFEB-GFP expressing plasmid. Transfected cells were subjected to live imaging of GFP and nuclei (using Hoechst 33342 staining). Left: representative images of cells with mostly cytosolic or mostly nuclear TFEB-GFP. Quantitative analysis of N = 3 experiments is shown on the right. Scale bar: 10 µm. (B) HEK293 cells expressing M5R-DREADD were starved of FBS or given fresh FBS-containing media for 2 hours before being treated with 1 µM CNO or 100 nM Torin1 for 32 minutes. After cellular fractionation, cytoplasmic (C) and nuclear (N) fractions were analysed by SDS-PAGE and western blotting. Quantitative analysis of N = 3 experiments is shown on the right. (C) HEK293 cells expressing M5R-DREADD were starved of FBS for 2 hours before stimulation with 1 µM CNO for 2 hours (where indicated). Lysosomes and nuclei were stained with 20 nM LysoTracker™ Red DND-99 and 2 µg/ml Hoechst 33342, respectively, before live imaging. Quantitative analysis of N = 3 experiments is shown on the right. Scale bar: 10 µm.

We then quantified lysosomes using a fluorescent lysosome-staining dye (Lysotracker). As shown in Figure 4C, serum starvation increased the intensity of Lysotracker signal and the number of Lysotracker-positive lysosomes, suggestive of enhanced lysosome biogenesis. This trend was not reversed by subsequent M5R stimulation by CNO, in contrast to the recovery in protein synthesis (Figure 3B). The fact that we did not observe a further increase of lysosomes suggests that M5R-driven dephosphorylation of TFEB is not sufficient to increase lysosome biogenesis in our experimental setting.

Taken together, M5R activation promoted protein synthesis without suppressing lysosome biogenesis. This observation was consistent with the differential effect of M5R stimulation on phosphorylation of S6K1 and 4EBP1, and that of TFEB (Figure 2D). We conclude that M5R regulates mTORC1 in a substrate/downstream-specific manner.

### Mechanistic insights into the M5R-mTORC1 signaling

We next addressed the mechanism of M5R-mTORC1 signaling. Because M5R activates the phospholipase C-protein kinase C (PKC) pathway downstream of G_q_ (Figure 5A, left) (10)(11), we examined the effect of the PKC inhibitor GF109203x. As shown in Figure 5B, CNO-triggered phosphorylation of S6K1 and 4EBP1 in M5R-DREADD cells was blocked by GF109203x (lanes 4 and 5, panels 1 and 3), suggesting that mTORC1 activation towards S6K1 and 4EBP1 requires PKC activity.

**Figure 5.**
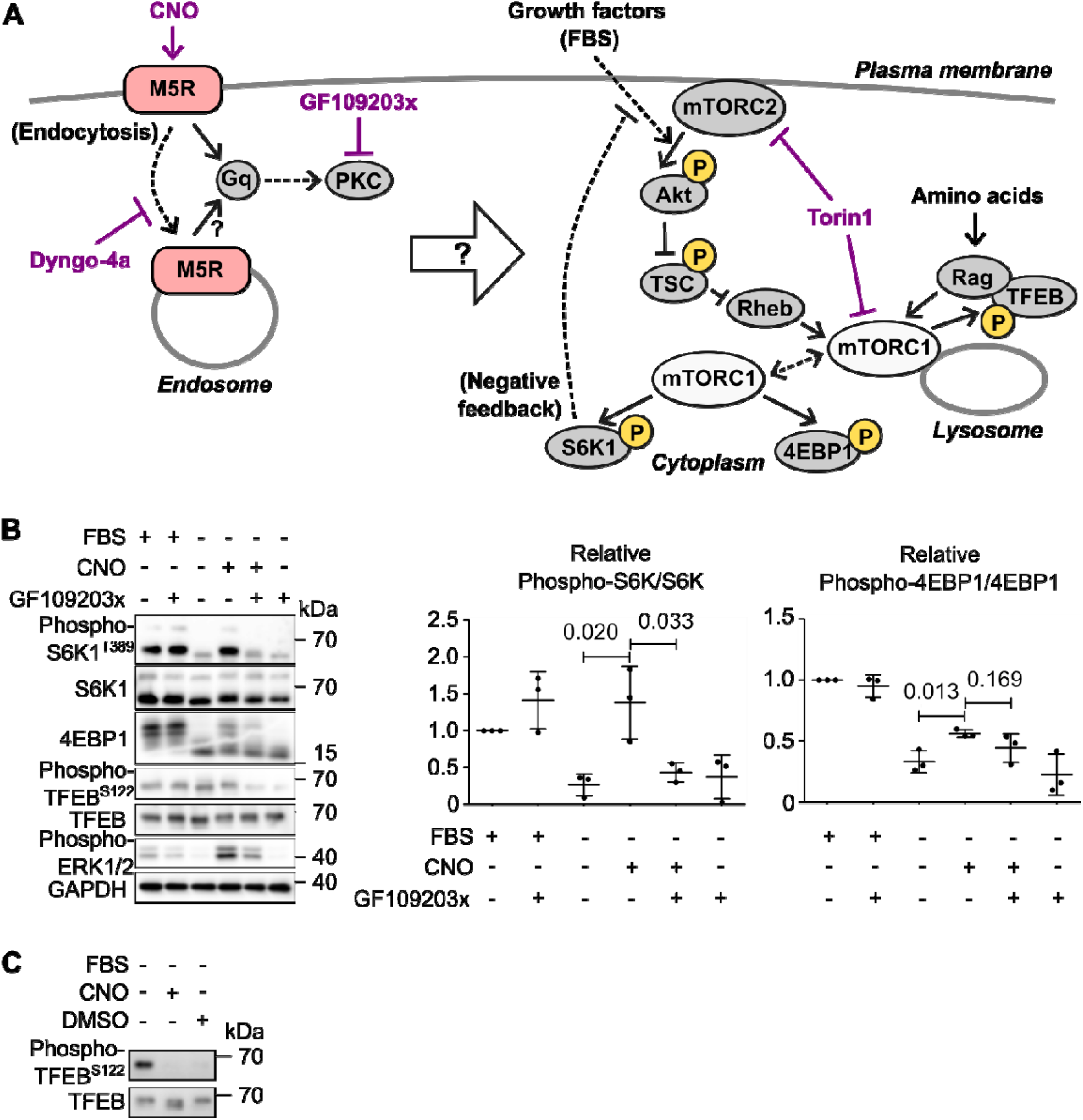
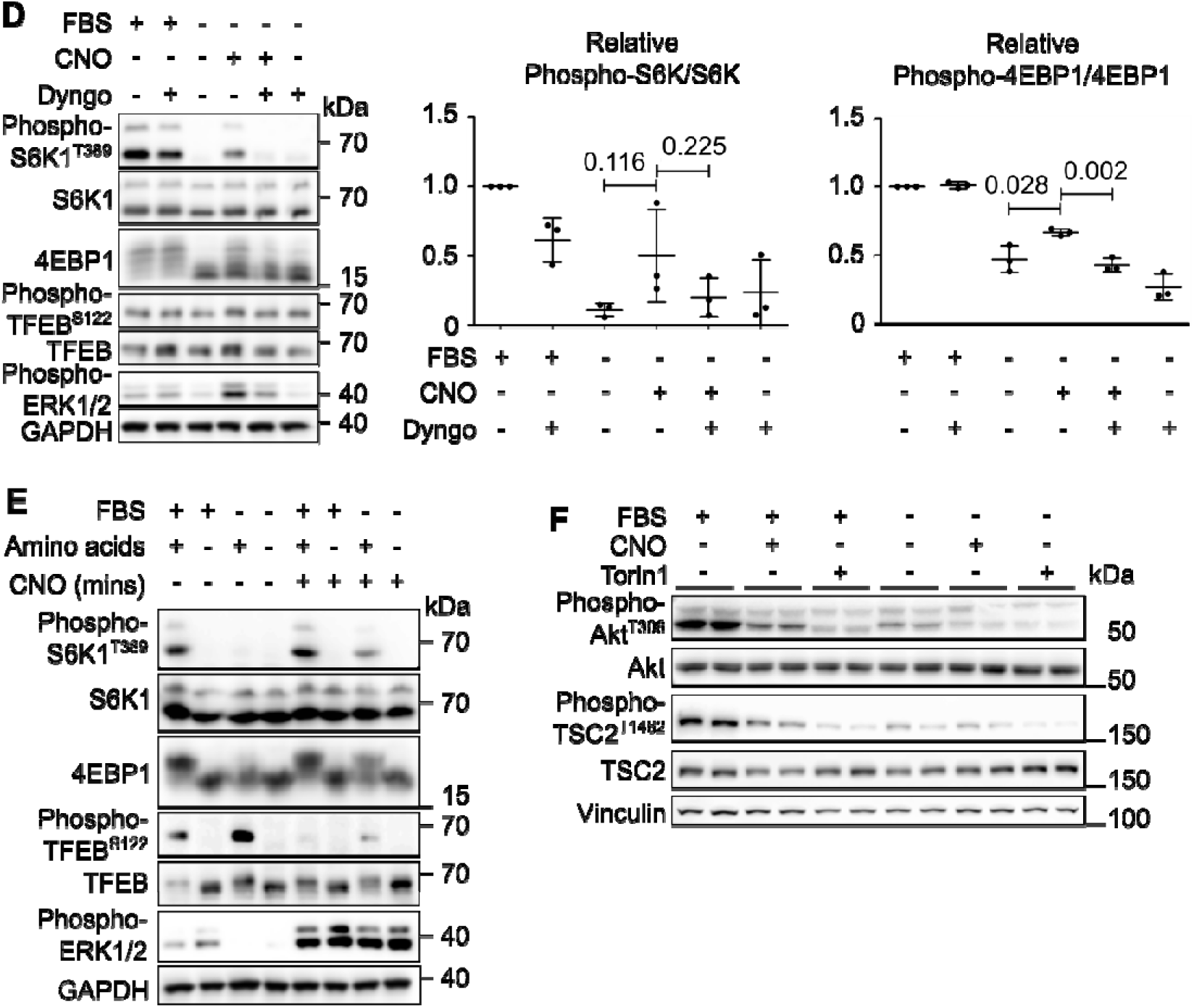
Mechanistic insights into the M5R-mTORC1 signaling. (A) Schematic outlying possible signaling modes of M5R to mTORC1. The points of action of CNO, GF109203x, Dyngo®-4a, and Torin1 are shown in purple. (B) HEK293 cells expressing M5R-DREADD were starved of FBS or given fresh FBS-containing media for 2 hours before being stimulated with 1 µM CNO for 32 minutes. PKC was inhibited using 5 µM GF109203x for 1.5 hours before CNO treatment. Cells were lysed and analysed by SDS-PAGE and western blotting. Quantitative analysis of N = 3 experiments is shown on the right. (C) HEK293 cells expressing M5R-DREADD were starved of FBS or given fresh FBS-containing media for 2 hours before being treated with 1 µM CNO for 32 minutes or DMSO at 1:2000 for 1.5 hours. Cells were lysed and analysed by SDS-PAGE and western blotting. (D) HEK293 cells expressing M5R-DREADD were starved of FBS or given fresh FBS-containing media for 2 hours before being stimulated with 1 µM CNO for 32 minutes. The Dynamin inhibitor Dyngo®-4a (to prevent endocytosis) was added at 10 µM for 30 minutes before CNO treatment. Cells were lysed and analysed by SDS-PAGE and western blotting. Quantitative analysis of N = 3 experiments is shown on the right. (E) HEK293 cells expressing M5R-DREADD were starved of FBS, amino acids, or both, or given fresh FBS-containing media for 2 hours before being stimulated with 1 µM CNO for 32 minutes. Cells were lysed and analysed by SDS-PAGE and western blotting. (F) HEK293 cells expressing M5R-DREADD were starved of FBS or given fresh FBS-containing media for 2 hours before being treated with 1 µM CNO or 100 nM Torin1 for 32 minutes. Cells were lysed and analysed by SDS-PAGE and western blotting.

We were unable to assess the role of PKC in the M5R-triggered dephosphorylation of TFEB, because TFEB did not respond to CNO in this experiment (lanes 3 and 4, panel 4). This was due to an unexpected effect of DMSO, which was used as the solvent of GF109203x as well as the mock control— we noticed that the DMSO treatment alone causes severe dephosphorylation of TFEB (Figure 5C). As an additional note, the effects of CNO and DMSO differed in that only the former caused a downshift of the total TFEB band (panel 2), suggesting that M5R also regulates posttranslational modification(s) of TFEB other than its Serine 122 phosphorylation.

As GPCRs signal from both the plasma membrane and endosomes (13), we asked where M5R signals to mTORC1. The remarkably slow kinetics of mTORC1’s response to CNO led us to hypothesize that M5R only signals to mTORC1 after being internalized. We addressed this possibility using the dynamin inhibitor Dyngo-4a that blocks endocytosis (19). As shown in Figure 5D (lanes 4 and 5, panels 1 and 3), Dyngo-4a significantly blocked phosphorylation of S6K1 and 4EBP1 triggered by M5R. This observation favours the post-internalization signaling model, as recently proposed by others (20). Our results did not inform the location where M5R signals to the mTORC1-TFEB axis, because of the issue of DMSO (Figure 5C), in which Dyngo-4a was dissolved.

mTORC1 is activated by the actions of Rag and Rheb small GTPases, which mediate amino acid and insulin/growth factor signals, respectively (Figure 5A, right). Notably, M5R failed to stimulate S6K1 and 4EBP1 phosphorylation in amino acid-starved cells (Figure 5E, lanes 2 and 6, panels 1 and 3; compare with serum-starved cells in lanes 3 and 7). This was not due to the failure of M5R activation, because ERK1/2 phosphorylation was similarly triggered by CNO in serum- and amino acid-starved cells (lanes 6 and 7, panel 6). Hence, M5R seems to activate the Rheb branch but not the Rag branch, the latter of which requires an amino acid signal.

Growth factors, including insulin, activate Rheb via the mTORC2-Akt cascade (Figure 5A, right). M5R does not activate this canonical upstream pathway, because CNO did not promote phosphorylation of Akt and TSC2, an Akt substrate (Figure 5F). These phosphorylation events were rather downregulated by CNO, which could be due to the known negative feedback regulation through S6K1 (Figure 5A, right).

Altogether, the most straightforward interpretation of our data is that M5R activates the mTORC1-S6K1/4EBP1 axis through the G_q_-PKC pathway after being internalized. This signal most likely impinges on Rheb, through unknown mechanisms independent from the canonical mTORC2-Akt-TSC pathway. How M5R inhibits the mTORC1-TFEB axis is less clear.

### mTORC1 regulation by FPR2 and β2AR

To gain broader insight, we asked whether substrate-specific regulation of mTORC1 is unique to M5R/G_q_-coupled GPCRs. We examined the G_i_-coupled formyl peptide receptor 2 (FPR2). FPR2-expressing HEK293 cells previously generated (21) were serum starved and treated with the peptide ligand WKYMVm, a potent FPR2 agonist. As shown in Figure 6A, WKYMVm triggered phosphorylation of S6K1 and 4EBP1 in 16 minutes (panels 1 and 3), though not affecting the cellular metabolic activity (Figure 6B). S6 phosphorylation was promoted as expected (Figure 6C). Unlike M5R, FPR2 did not strongly inhibit TFEB phosphorylation (Figure 6A, panel 4). Lysosome biogenesis was unaffected (Figure 6D).

**Figure 6.**
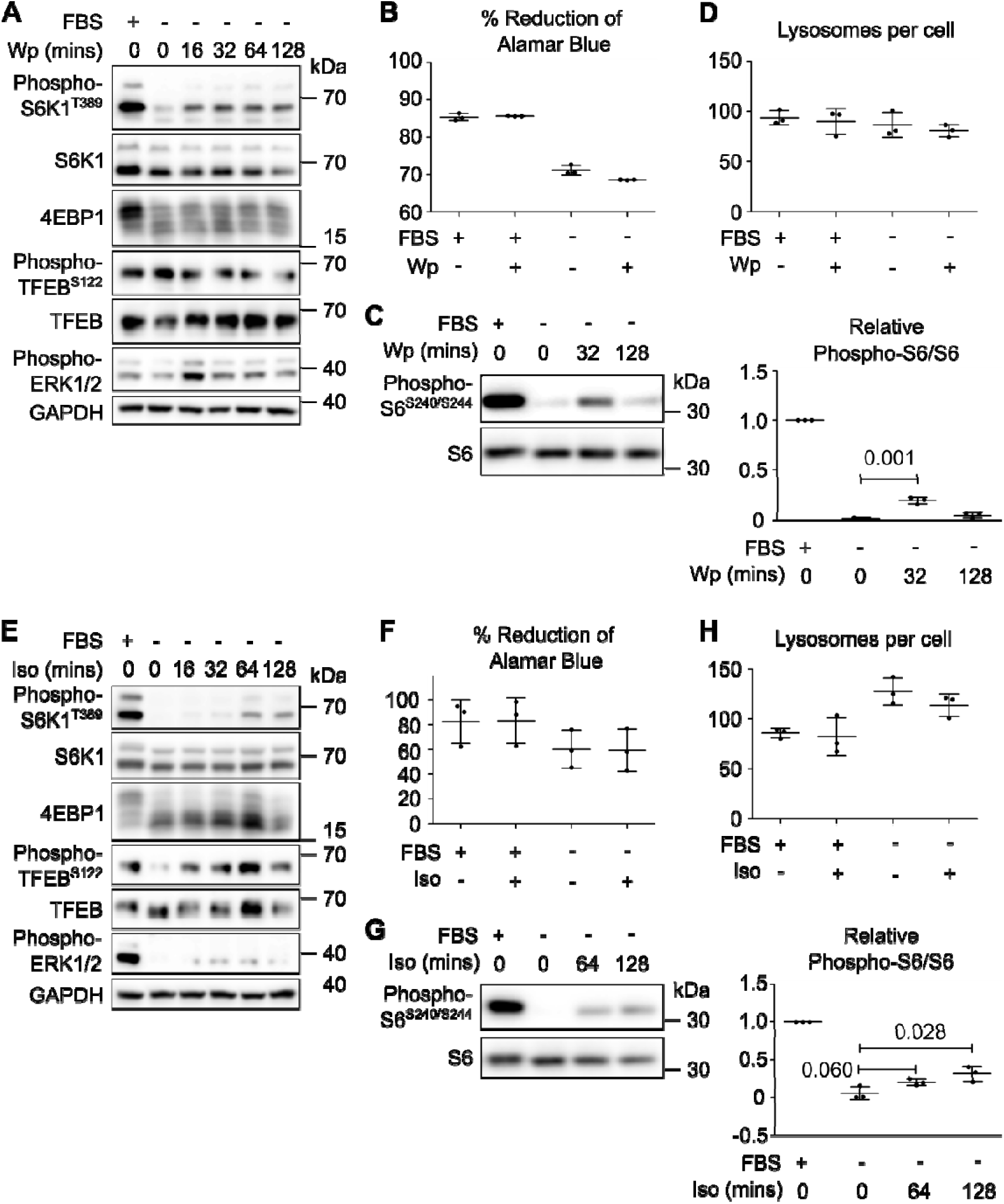
Differential response of mTORC1 substrates to other GPCRs. (A) HEK293 cells expressing FPR2 were starved of FBS or given fresh FBS-containing media for 2 hours before being stimulated with 1 µM WKYMVm peptide (Wp) for up to 128 minutes. Cells were lysed and analysed by SDS-PAGE and western blotting. (B) Metabolic activity of HEK293 cells expressing FPR2 was measured by the reduction of Alamar Blue after overnight stimulation with 1 µM WKYMVm peptide with or without serum starvation. (C) HEK293 cells expressing FPR2 were starved of FBS or given fresh FBS-containing media for 2 hours before being stimulated with 1 µM WKYMVm peptide for the indicated times. Cells were lysed and analysed by SDS-PAGE and western blotting. Quantitative analysis of N = 3 experiments is shown on the right. (D) HEK293 cells expressing FPR2 were starved of FBS for 2 hours before stimulation with 1 µM WKYMVm peptide for 2 hours (where indicated). Lysosomes and nuclei were stained with 20 nM LysoTracker™ Red DND-99 and Hoechst 33342 before live imaging. The result of a quantitative analysis of N = 3 experiments is shown. (E) Wild-type HEK293 cells were starved of FBS or given fresh FBS-containing media for 2 hours before being stimulated with 1 µM isoprenaline (Iso) for up to 128 minutes. Cells were lysed and analysed by SDS-PAGE and western blotting. (F) Metabolic activity of wild-type HEK293 cells was measured by the reduction of Alamar Blue after overnight stimulation with 1 µM isoprenaline with or without serum starvation. (G) Wild-type HEK293 cells were starved of FBS or given fresh FBS-containing media for 2 hours before being stimulated with 1 µM isoprenaline for the indicated times. Cells were lysed and analysed by SDS-PAGE and western blotting. Quantitative analysis of N = 3 experiments is shown on the right. (H) Wild-type HEK293 cells were starved of FBS for 2 hours before stimulation with 1 µM isoprenaline for 2 hours (where indicated). Lysosomes and nuclei were stained with 20 nM LysoTracker™ Red DND-99 and Hoechst 33342 before live imaging. The result of a quantitative analysis of N = 3 experiments is shown.

We then examined G_S_-coupled GPCRs. We treated wild-type HEK293 cells, which endogenously express the G_S_-coupled beta-2 and −1 adrenergic receptors (β2AR and β1AR), with their ligand isoprenaline. Isoprenaline triggered phosphorylation of S6K1 in 64 minutes (Figure 6E, panel 1), but did not cause a clear band shift of 4EBP1 (panel 3). Isoprenaline did not affect cellular metabolic activity (Figure 6F), despite increased S6 phosphorylation (Figure 6G). TFEB phosphorylation was not inhibited (Figure 6E, panel 4), with lysosome biogenesis unaffected (Figure 6H).

To summarize, M5R, FPR2, and β2AR/β1AR all activated mTORC1 towards S6K1. M5R and FPR2 also activated the mTORC1-4EBP1 branch. However, M5R inhibited mTORC1 towards TFEB, uniquely among the three GPCRs. Hence, different GPCRs regulate mTORC1 with varied substrate specificities.

## Discussion

This study uncovered the substrate/downstream-specific mode of mTORC1 regulation by GPCRs, revealing an additional layer of complexity to this signaling axis. Notably, the response of TFEB was very different (even opposite for M5R) from that of canonical mTORC1 substrates, S6K1 and 4EBP1. A lesson to be drawn from this is that we need to monitor multiple mTORC1 substrates including TFEB when assessing the effects of GPCRs. This perspective has been largely overlooked in previous studies, which have predominantly focused on canonical mTORC1 targets (14).

Mechanistically, the unique response of TFEB upon GPCR stimulation can be partly explained if GPCRs signal to mTORC1 through the Rheb branch, which TFEB is uniquely unresponsive to. This interpretation aligns with our observation that M5R stimulation overrides FBS starvation but not amino acid starvation (Figure 5E). It is indeed known that the MEK-ERK1/2 cascade, which acts downstream of many GPCRs including the ones examined here, activates Rheb by inhibiting its negative regulator TSC complex (22). Another factor could be the subcellular location of phosphorylation events. Given that TFEB is phosphorylated at lysosomes rather than the cytoplasm as proposed for S6K1 and 4EBP1 (5), GPCRs may exert substrate-specific effects by modulating mTORC1 localization or location-specific activities. The involvement of phosphatases antagonizing mTORC1 should also be considered, which could be different for each mTORC1 substrate (4).

The non-canonical response of TFEB observed in this and other studies indicates the regulatory decoupling of a catabolic process (lysosome biogenesis) from an anabolic process (protein synthesis), even though both are governed by mTORC1. A similar decoupling is observed in the budding yeast model, in which protein synthesis and autophagy are regulated by spatially distinct TORC1 pools (23). This may look counterintuitive from the traditional view in the TORC1 field that activation of anabolism or catabolism should coincide with inhibition of the other. Such coupling would make sense in the context of nutrient response, where cells adapt their bulk mass to nutrient availability. However, the situation is different when cells respond to other stimuli such as growth factors or stresses. Cells would need to change themselves qualitatively by remodelling the proteome, not necessarily changing their total mass. For that, they would need to synthesize proteins that are needed in the new environment, and simultaneously degrade proteins not needed anymore. The latter catabolic process and subsequent recycling of resources (e.g., amino acids) fuel the former anabolic processes. In such a situation, it would be beneficial to activate both anabolism and catabolism by decoupling their regulations.

There is another potential benefit of metabolic decoupling in light of cell proliferation. M5R robustly stimulated cellular metabolic activity (Figure 1D). Under this condition where cells actively proliferate, it would be beneficial to keep TFEB dephosphorylated and generate lysosomes needed for daughter cells. TFEB dephosphorylation was indeed not observed upon FPR2 or β2AR/β1AR activation, where cellular metabolic activity was unaffected (Figure 6).

The supreme druggability of GPCRs, and their tissue/context-specific expression patterns, make GPCR ligands attractive therapeutic tools to manipulate mTORC1 pathway to treat cancer, diabetes, neurodegeneration, and aging. The substrate/downstream specificity revealed in this study further enhances the potential of GPCR ligands, as it theoretically enables selective manipulation of the desired downstream functions of mTORC1, which should be more effective and safer than pan-mTORC1 manipulation. This transformative potential warrants a comprehensive investigation of the mTORC1 substrate specificity for a wider range of GPCRs.

## Materials and methods

### Antibodies and reagents

All antibodies were purchased from Cell Signaling Technologies except anti-GAPDH (Proteintech 60004-1-Ig). Anti-S6K (2708), anti-phospho-S6K Thr389 (9205), anti-4EBP1 (9644), anti-TFEB (4240), anti-phospho-TFEB Ser122 (87932), anti-S6 (2217), anti-phospho-S6 Ser240/Ser244 (5364) anti-phospho-p44/42 MAPK (Erk1/2) Thr202/Tyr204 (4370), anti-anti-Vinculin (13901), anti-Lamin A/C (4777), anti-Akt (4691), anti-phospho-Akt Ser473 (4060), anti-TSC2 (4308), and anti-phospho-TSC2 Thr1462 (3617) were used at 1:1000. Anti-GAPDH was used at 1:30,000. Unless otherwise stated, common reagents were purchased from Thermo Fisher Scientific.

### Cell culture

All cell lines (HEK293 and derivatives) were cultured in DMEM high glucose (Merck D5671) supplemented with 1 mM sodium pyruvate (Thermo Fisher Scientific 11360-039), 1% glutamine (Merck G7513) and 10% FBS (Biosera 10011500) at 37°C in an atmosphere of 5% CO_2_. Cell lines were regularly tested for the presence of mycoplasma contamination. FPR2 stably expressing cells were previously generated as described in (21). DMEM without amino acids (VWR USBID9800-13) was supplemented with glucose to match DMEM high glucose above, 1 mM sodium pyruvate and 10% dialysed FBS (lacking amino acids). For dialysing FBS, the same FBS batch used for all experiments was dialysed against PBS twice overnight at 4°C using a 3500 Da cutoff Snakeskin™ dialysis membrane (Thermo Fisher Scientific 88242).

### Muscarinic Acetylcholine Receptor M5 DREADD stable cell line

HEK293 cells were from the ATCC. The M5 DREADD receptor was generated based on the M3R DREADD (24). Briefly, M5R cDNA was obtained from cDNA.org and subcloned into the Flag pcDNA3.1+ zeo (21). Site-directed mutagenesis (QuickChange, Agilent) was then used to introduce the mutations in the 3^rd^ and 5^th^ transmembrane domain (Y111C and A201G) to confer sensitivity to CNO. Stables were generated by selection with zeocin and identified by anti-Flag immunofluorescence determined by microscopy.

### GPCR stimulation

All cell lines were washed with PBS and then starved of amino acids or FBS for 2 hours before treatment with the GPCR antagonists at 1 µM: CNO (Tocris/bio-techne 6329), Isoprenaline (Sigma/Merck I6504) or WKYMVm (Tocris/BioTechne 1800) in M5R-DREADD, wild-type or FPR2-expressing HEK293 cells, respectively, for the indicated timepoints. For inhibitor experiments: 10 µM Dyngo®-4a (Abcam ab120689) or 5 µM PKC inhibitor GF109203x (Sigma/Merck B6292) was added to the culture media 30 minutes or 1.5 hours, respectively, before GPCR stimulation. 100 nM Torin1 was applied for 32 minutes.

### Global translation assay

Cells were stimulated as above, and puromycin (Thermo Fisher Scientific 227420500) was added 15 minutes before lysis at 5 µg/ml as per the SUNSET assay protocol (17).

### Cellular fractionation

Cellular fractionation was performed with the NE-PER™ Nuclear and Cytoplasmic Extraction Reagents kit (Thermo Fisher Scientific, 78835). The cells were seeded in 6-well plates at a cell density of 4x10^5^ cells/well. The next day they were treated as specified in the figure legends. After treatment, the cells were washed once with PBS (Substrate and Sterile Centre, University of Copenhagen). The washed cells were lysed with the cytosolic extraction reagents I and II (CERI and CERII) provided in the kit, to induce disruption of the cell membranes and the release of cytosolic proteins. After being scrapped in cold CERI buffer containing protease and phosphatase inhibitors (500 μM phenylmethylsulfonyl fluoride (PMSF [Sigma, P7626]), 1 mM benzamidine hydrochloride [Sigma, 199001], 1 μg/ml leupeptin [Sigma, L2884], 1 μg/ml pepstatin A [Sigma, P5318], 1 μM microcystin-LR [Enzo Life Sciences, ALX-350-012], 1 mM sodium orthovanadate [Sigma, S6508]), the cells in CERI buffer were dispersed by brief vortexing, incubated on ice for 10 min and then, mixed with CERII buffer and further incubated on ice for 1 min. The separation of cytosolic proteins from the nuclear pellet was performed by centrifugation at 16,000 x g for 5 min at 4°C. The supernatant containing the cytosolic protein fraction was collected in a new tube and denatured at 100°C for 5 min in Laemmli sample buffer (50 mM Tris, pH 6.8, 2% sodium dodecyl sulfate [Sigma, 74255], 0.5 mM EDTA (Sigma, E5134), 10% glycerol [Sigma, G5516], 0.01% bromophenol blue [Sigma, 114391]). The pellet containing the nuclei was washed three times with cold PBS. Nuclear protein fractions were obtained by extraction from the nuclear pellet with Laemmli sample buffer at 100°C for 7 min. The denatured cytosolic and nuclear protein fractions were stored at −20°C until Western blot analysis.

### Western blotting

Cells were washed with PBS before being lysed directly in high SDS lysis buffer (3% SDS, 60 mM sucrose, 65 mM Tris-HCl pH 6.8), denatured at 95°C for 5 minutes, homogenised by sonication and cleared by centrifugation at 18,000 x g for 30 minutes. Lysates were quantified by a linearised Bradford assay (Abcam ab119216) using a 595 nm/450 nm ratio.

25 µg of proteins were separated in SDS-PAGE gels before being transferred to nitrocellulose membranes (VWR 10600002). After ponceau staining, membranes were blocked with 5% milk/TBST (for anti-TFEB, phospho-TFEB, GAPDH, and phospho-MAPK antibodies), 3% milk/TBST (for anti-Vinculin, Lamin A/C, Akt, phospho-Akt, TSC2, and phospho-TSC2 antibodies) or 5% BSA/TBST (for anti-S6K, phospho-S6K, and 4EBP1 antibodies) and probed with antibodies overnight at 4°C in 5% BSA/TBST (for anti-S6K, phospho-S6K, 4EBP1, TFEB, phospho-TFEB, and phospho-MAPK antibodies), 3% milk/TBST (for anti-Vinculin, Lamin A/C, Akt, phospho-Akt, TSC2, and phospho-TSC2 antibodies) or 5% Milk/TBST (for anti-GAPDH antibody). Membranes were visualised after secondary antibody incubation (Cell Signaling Technologies 7074 and 7076) and washing with TBST using ECL substrate (Thermo Fisher Scientific 12994780). Membranes were imaged in a Gbox gel/blot imaging system (Syngene) and densitometry analysis was performed using ImageJ (25).

### TFEB-GFP microscopy

HEK293 M5R-DREADD cells were transfected overnight using Fugene® HD (Promega E2311) with TFEB-GFP (MRC PPU Reagents and Services DU67822). Transfected cells were re-seeded onto 8 well chamber slides (Thistle Scientific IB-80826) and allowed to recover overnight. Wells were washed with PBS and starved of FBS for 2 hours except for AA starvation control wells which were only starved of AA for a total of 45 minutes before imaging. Cells were stained with 2 μg/ml Hoechst 33342 (Thermo Fisher Scientific 62249) and stimulated with 1 µM CNO for 32 minutes before imaging. Live cell images were acquired on an Ultraview VoX spinning disk confocal microscope (PerkinElmer) with a 40x objective. 6 full fields of view for each condition were captured per experiment, and the percentage of the total TFEB-GFP signal that was nuclear was then measured. Images were analysed with ImageJ, defining GFP signals overlapping with Hoechst signals as nuclear TFEB-GFP.

### Lysosome microscopy

Cells were seeded overnight onto 8 well chamber slides and allowed to recover overnight. Cells were then washed with PBS and starved of FBS for 2 hours followed by 2 hours of GPCR stimulation (see above). 20 nM LysoTracker™ Red DND-99 (Thermo Fisher Scientific L7528) and 2 ug ml^−1^ Hoechst 33342 were added to the wells for 30 minutes before imaging. Live images were acquired on an Ultraview VoX spinning disk confocal microscope (PerkinElmer) with a 63x objective. 6 full fields of view for each condition were captured per experiment, and the numbers of lysosomes per cell were measured. Images were analysed using the find maxima function in ImageJ.

### Alamar Blue assay

Cells were seeded overnight in 96 well plates before being washed with PBS and treated with DMEM +/− FBS and +/− GPCR agonist (see above) for 18 hours. Media was then replaced with DMEM + 10% Alamar blue solution (Bio-Rad BUF012A) for 3 hours before absorbance at 570 nm and 600 nm was measured on a Clariostar plus plate reader (BMG Labtech). Analysis was performed as per manufacturer’s recommendations.

## Abbreviations

mTORC1: mammalian/mechanistic Target of Rapamycin Complex 1
GPCR: G protein-coupled receptor
M5R: muscarinic acetylcholine receptor M5
FPR2: formyl peptide receptor 2
β2AR: beta-2 adrenergic receptor
DREADD: designer receptor exclusively activated by designer drugs
CNO: Clozapine N-oxide
PKC: protein kinase C

## Acknowledgments

We thank Fiona Murray (University of Aberdeen) for helpful suggestions; Colin Ferguson and the Microscopy and Histology Core Facility members Lucinda Wight and Debbie Wilkinson (University of Aberdeen) for technical support. This work was funded by BBSRC (BB/V016334/1 and BB/X018229/1 to RH), British Heart Foundation (PG/19/30/34327 to DT), Novo Nordisk Foundation (NNF23SA0084103 and NNF24OC0092181 to KS), and the University of Aberdeen. The authors declare no conflict of interest.

## Author contributions

Conceptualization: SJA, DT, JH, and RH; Methodology: SJA, FN, WR, KT, and JH; Investigation: SJA, FN, WR, KT, and JH; Formal Analysis: SJA, FN, and JH; Validation: SJA, FN, WR, and JH; Supervision: KS, DT, JH, and RH; Funding acquisition: KS, DT, JH and RH; Visualization: SJA, JH, and RH; Writing – original draft: SJA, FN, and RH; Writing – review & editing: SJA, KS, DT, JH, and RH.

